# Crowdsourced riboregulators reveal design principles for programmable RNA switching

**DOI:** 10.64898/2026.07.08.737257

**Authors:** James M. Robson, Gabrielle Moussas, Dayna Francis, Alexander A. Green

## Abstract

RNA-based sensors offer powerful and programmable control of gene expression, yet our understanding of the structural principles that govern their potential design space remains incomplete. Here, we challenged a community of designers to generate novel riboregulators capable of activating translation in response to specific RNA targets. Participants submitted diverse sequence architectures, which were synthesized and evaluated in a cell-free transcription-translation system. Across 100 designs, community-generated riboregulators displayed wide variability in activation dynamics, fold change, and structural features, outperforming some canonical toehold-switch designs and achieving up to 80-fold activation. Structural ensemble analyses identified accessibility patterns near the ribosome binding site that distinguish high- from low-performing regulators, highlighting the central role of RBS sequestration and release in modulating expression. Together, we demonstrate community-driven design can expand the accessible structural space of riboregulators and uncover mechanistic features governing translational activation. Our findings establish quantitative links between RNA folding energetics and gene expression output, providing design principles for next-generation programmable RNA sensors.

## INTRODUCTION

RNA-based sensors enable programmable regulation of gene expression by coupling recognition of diverse molecular inputs to structural rearrangement. Across cellular and cell-free systems, naturally occurring and engineered RNA regulators control transcription^1–4^, translation^5,6^, RNA stability^7^, and splicing^8^ through conformational switching. Major classes of RNA sensors include transcriptional riboswitches^9^, small RNA regulators^10^, CRISPR-associated regulation^11^, and translational riboregulators^12,13^. Due to the unique advantages of RNA as a highly programmable substrate, RNA sensors have been deployed in molecular diagnostics^14–17^, gene circuit construction^18,19^, and programmable therapeutic strategies^20,21^.

Among engineered RNA sensors, toehold switches represent one of the most versatile and modular scaffolds for regulating translation^6^. Toehold switches sequester the ribosome binding site (RBS) and start codon within a hairpin structure, preventing translation in the absence of a cognate target RNA. Upon target hybridization to a toehold domain, strand displacement unfolds the hairpin, exposing the RBS and enabling translation. The programmability of toehold switches has allowed them to be adopted in electrochemical and paper-based detection platforms^22–25^ and gene control in both cellular and cell-free systems^13^.

Despite these advancements, the design of high-performance riboregulators remains challenging. Translation initiation is governed not only by sequence complementarity between target and switch, but also by RNA structural dynamics, ribosome binding energetics, and kinetic accessibility^26–28^. Translation efficiency depends on the free energy landscape of RBS accessibility, including contributions from mRNA secondary structure, ribosomal standby site interactions, and unfolding energetics^29,30^. Thermodynamic stability alone does not determine regulatory output, and therefore designing riboregulators with only thermodynamic folding software has resulted in many computationally designed riboregulators exhibiting unpredictable activation levels, incomplete repression, or reduced dynamic range^18^. Translational control emerges from the interplay between sequence, structure, kinetics, and ribosome engagement.

Machine learning and deep learning techniques have overcome some of these limitations and revolutionized the prediction of RNA sequence-structure-function relationships^31–35^. By training on high-throughput datasets, computational frameworks decode the rules governing sequence and structural robustness across thousands of variants. However, the theoretical design space for RNA riboregulators spans not only the sequence space of the target interaction domain (typically 4^30^ or ∼ 10^18^ for a 30-nucleotide RNA target sequence), but also the combinatorial possibilities of structural space. When considering the possibilities of stem lengths, loop configurations, mismatches, bulges, multibranch junctions, and alternative RBS sequestration geometries, even the largest experimental libraries represent only a small fraction of the total design landscape^6^. Consequently, most design strategies are constrained by assumptions about optimal RNA sensor folding and potentially overlook unconventional structural solutions.

Crowdsourcing offers a complementary strategy for navigating large combinatorial design landscapes^36–41^. Citizen-science initiatives such as Foldit^42^, Eterna^43^, and the platform Folding@home^44^ have demonstrated that distributed problem-solving can uncover unexpected structural designs, outperform automated algorithms, and accelerate scientific discovery. In molecular design contexts, crowdsourced science has explored unconventional structural motifs that feature designs beyond fixed optimization rules^36^. The success of community-driven approaches suggests that crowdsourced design may be particularly well suited for riboregulators, where sequence-structure relationships are governed by predictable base pairing.

Here, we employ a community-driven design strategy to explore the structural landscape of translational riboregulators beyond the canonical toehold-switch geometry. By challenging a network of designers to engineer RNA switches, we sampled a highly diverse array of sequence-structure architectures. Designs were characterized using cell-free transcription-translation assays, allowing for quantitative mapping of regulatory performance to RNA folding energetics and translation initiation kinetics. Our findings reveal structural principles that drive high-performance translational repression and activation and demonstrate that unconventional architectures, identified through community-guided exploration, can significantly expand our understanding of programmable RNA sensing.

## RESULTS

To explore the structural design space of translational RNA regulators, we developed a crowdsourced riboregulator framework (Figure 1A, B). A community was challenged to design RNA sensors capable of activating translation in the presence of defined RNA targets, using standard thermodynamic modeling packages to generate sequences that would fold differentially with a target RNA. Participants were provided with a standard RBS sequence, but no additional constraints were given on tools that the community could use to generate their designs. We solicited 100 total designs (Supplementary Figure 1, Supplementary Table 1), assembled the regulators into the 5’ untranslated region (UTR) of a GFPmut3b reporter, and characterized the switching behavior in a cell-free transcription-translation reaction with or without cognate target RNA (Figure 1 C, D). Fluorescence measurements revealed diverse activation dynamics across the library (Figure 1E, F). While many designs remained weakly responsive and demonstrated no activation, a subset exhibited rapid activation with half-maximal response times near two hours, achieving maximal GFP fluorescence with respect to positive controls. We then quantified the fold change of each crowdsourced sensor using two metrics: rate and fluorescence (Figure 1G). We found a wide distribution of riboregulator activation, with some sensors exceeding 80-fold change in the rate of GFP production.

**Figure 1.**
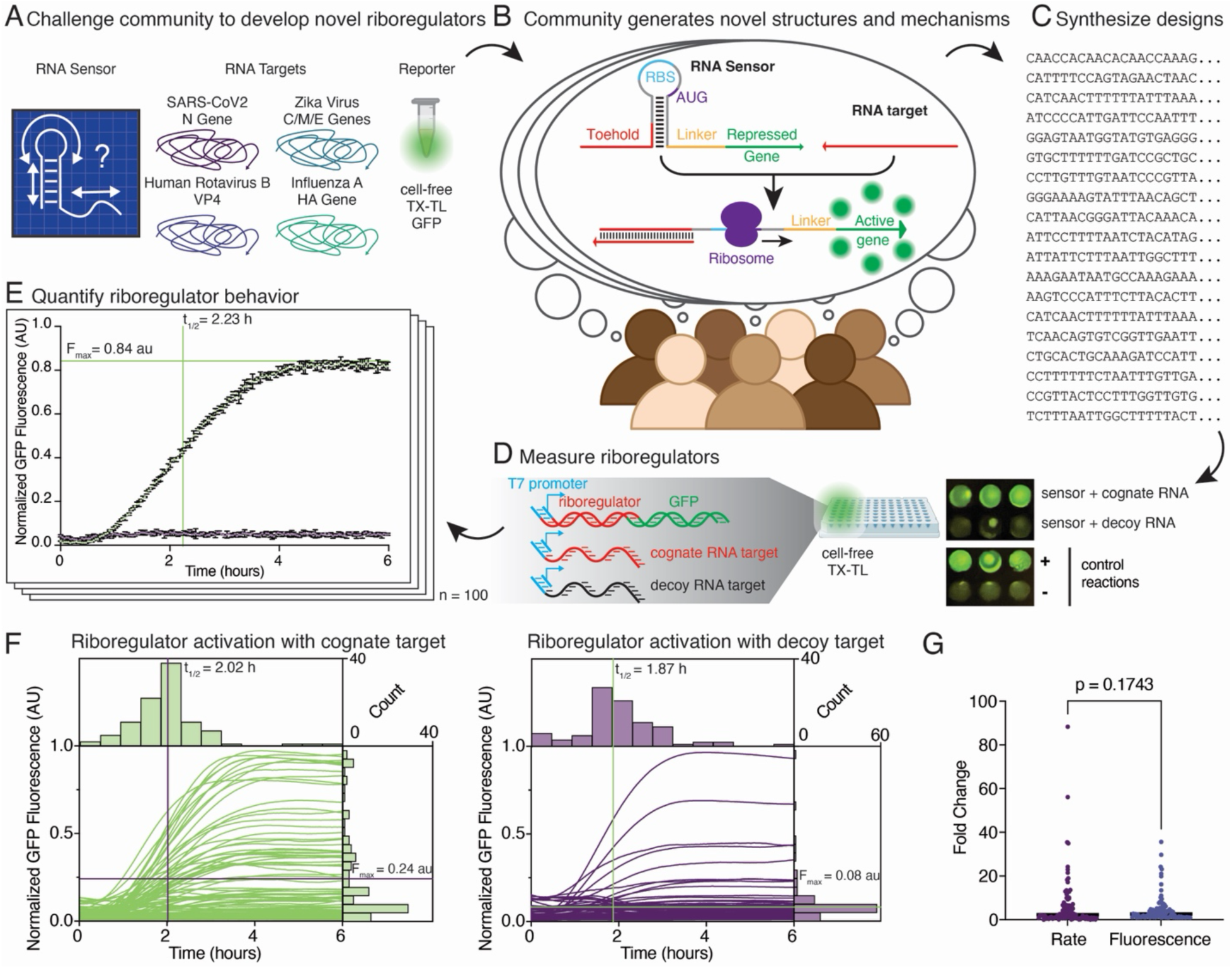
Characterization of crowdsourced translational riboregulators. **(A)** Participants were tasked with designing riboregulators responsive to defined RNA targets, including fragments from the SARS-CoV-2 nucleocapsid (N) gene, zika virus capsid/precursor membrane/envelope (CME) genes, human rotavirus outer capsid VP4 gene, and influenza A hemagglutinin (HA) gene. **(B)** Crowdsourced designs were required to embed a ribosome binding site (RBS) and start codon within structural elements upstream of GFP. Target RNA binding induces structural rearrangement and translational activation. **(C)** Community designs were synthesized as DNA oligonucleotides and assembled downstream of a T7 promoter and in the 5’ untranslated region of GFP. **(D)** Riboregulator constructs were evaluated in a cell-free transcription-translation (TX-TL) system under cognate target and decoy target conditions. **(E)** Representative GFP fluorescence trajectory showing activation dynamic for sensor 93, including F_max_ and t_1/2 max_, the average maximal GFP fluorescence intensity and average time to half maximal fluorescence, respectively. **(F)** GFP fluorescence trajectories for 100 crowdsourced riboregulators. Each line represents the smoothed average of n=3 biological replicates per sensor. **(G)** Fold-change activation between cognate and decoy target across the 100-design library (p=0.1743, two-tailed t-test). Bars represent the mean.

Community solicited designs were compared with the canonical toehold switch architecture using the versatile in-silico targeting analysis (VISTA)-designed toehold switch (Supplementary Figure 2, Supplementary Table 2). Analysis of the OFF-state leakage revealed that the crowdsourced designs exhibited significantly different baseline fluorescence levels compared to VISTA, though both remained relatively low (Supplementary Figure 2B). A more pronounced disparity was observed in the ON-state performance: the VISTA designs achieved consistently higher normalized GFP fluorescence compared to the more variable distribution of the crowdsourced designs (Supplementary Figure 2C). The crowdsourced designs that did successfully activate demonstrated significantly faster switching kinetics, with a lower mean time to half-maximal fluorescence (t_1/2max_) compared to the VISTA-designed controls. However, The VISTA designs achieved a significantly higher mean fluorescence fold change compared to the crowdsourced designs (Supplementary Figure 2E). Similarly, the fold change in rate of fluorescence was significantly greater in the VISTA baseline than the crowdsourced population. While the crowdsourced designs offer a trade-off in maximal GFP fluorescence intensity and leaky off state, the results suggest they provide a potential advantage in the speed of translational response to target activation.

To better understand why crowdsourced designs featured enhanced translational response upon switching, we first quantitatively assessed key structural features from the crowdsourced library (Figure 2A, B; Supplementary Table 3). Community submissions spanned a wide structural range. Sensor lengths varied substantially, as did the positioning of target-binding domains relative to the transcription start site (TSS) and RBS (Supplementary Figure 3A). Individual sensor domains ranged in MFE (ΔG_sens_) from -1.2 to -73.5 kcal/mol, but viral target regions selected by the community featured strikingly little structure, with an average MFE (ΔG_targ_) of -1.5 kcal/mol and high enrichment in A-U bases (Supplementary Figure 3B).

**Figure 2.**
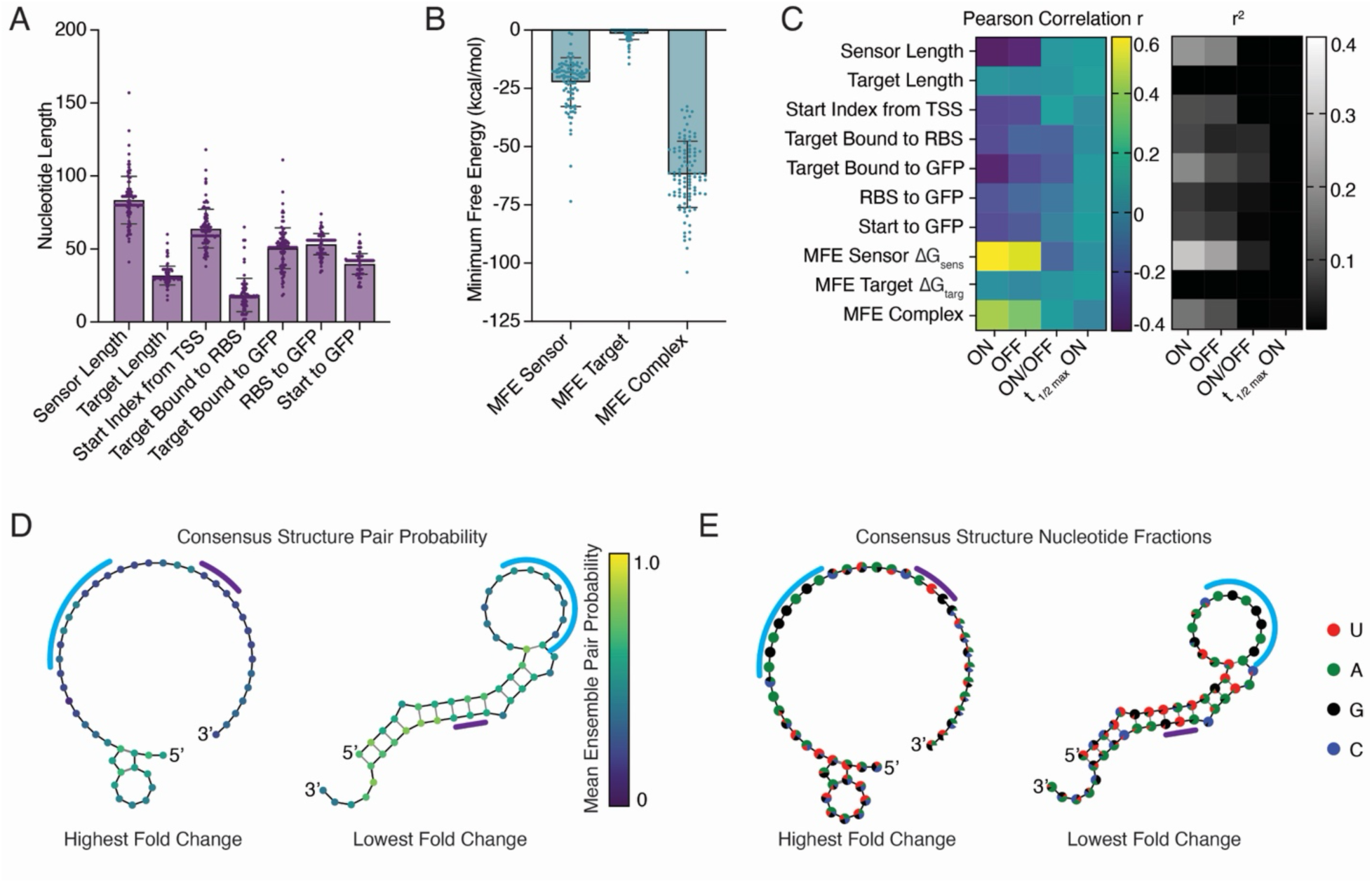
Structural and energetic diversity across community-generated riboregulators. **(A)** Variation in sensor length and target-binding domain position relative to the transcription start site (TSS) and RBS, and distance between target base pairing and conserved sequence features across the library. **(B)** Minimum free energy (MFE) distributions for sensor, target, and sensor-target complexes highlighting the wide variability in MFE for sensors and relatively unstructured targets chosen by the community. **(C)** Pearson correlation matrix linking structural and sequence features to functional outputs (ON fluorescence, OFF fluorescence, fluorescence fold change, and activation kinetics). **(D)** Consensus ensemble pairing probability comparison between highest- and lowest-performing (n=10) riboregulators, revealing distinct accessibility patterns near the RBS and translation initiation region. **(E)** Nucleotide frequency distributions across consensus structures for high- vs low-performing designs, indicating compositional biases toward A-U bases associated with functional performance. Bars represent mean and whiskers indicate the standard deviation (SD).

To connect structural features with functional output, we computed Pearson correlations between structural/energetic metrics and experimental phenotypes (Figure 2C). Sensor MFE exhibited the strongest correlation with ON state fluorescence (Pearson r = 0.54) indicating that intrinsic sensor structure influences maximal translational output. Several architectural features like the length of the sensor and distance between the start codon and TSS showed weaker negative correlations, indicating that longer 5’ UTRs reduce maximal fluorescence activation. Ensemble-level structural comparisons between the highest- and lowest-performing designs revealed distinct pairing probability distributions (Figure 2D). Consensus secondary structures were calculated using a majority-rule approach to determine the most frequent structural state—either paired or unpaired—at each aligned position across a population of individual RNA secondary structures (see Consensus Structure Calculation in Methods). High fold-change sensors displayed largely unstructured RBS sequestration within 20 nucleotides of the RBS, whereas low-performing designs exhibited significant pairing within the translation initiation region. Nucleotide composition analysis across the consensus structure (Figure 2E) further highlighted subtle nucleotide biases associated with high-performance architectures. However, given the composition of the highest three crowdsourced sensors (Supplementary Figure 3C), higher thermodynamic stability (more negative ΔG_sens_) does not always correlate with the highest fold change. For example, sensor 93 exhibits the highest fold change despite having a moderate ΔG_sens_ of -14.2 kcal/mol. While a stable OFF state is necessary, both the energetics and kinetics of opening are likely key factors deciding peak performance.

We next examined the relationship between RNA folding energetics and protein expression. Given the central role of ribosome accessibility in determining translation initiation^26–29^, we sought to evaluate the biophysical drivers of sensor performance. In existing thermodynamic models of ribosome binding, the translation initiation rate is determined by the energetic penalty required to transition the RBS to an accessible state suitable for ribosome loading^26^. The translation initiation rate (*k_init_*) scales exponentially with the total ribosome-mRNA binding free energy (ΔG_total_), such that:

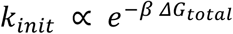

where *β* is the Boltzmann constant. Translation initiation therefore depends on the energetic balance between favorable ribosome-mRNA interactions and the penalty required to unfold inhibitory mRNA structures overlapping the ribosome footprint.

Consistent with this model, weakly structured RBS regions facilitate ribosome loading, whereas stable secondary structures impose energetic penalties that reduce translation initiation efficiency (Figure 3A)^29^. Correlation analysis confirmed that structural accessibility within the local RBS environment is a key driver of protein expression output (Figure 3B, Supplementary Table 4). We observed strong associations between the free energy of regions surrounding the RBS and experimental fluorescence across the library, particularly the MFE between the target binding domain and start of GFP (ΔG_RBS_, Pearson r = 0.64). MFE values calculated for windows surrounding the RBS (ranging from ±5 to ±30 nt) showed increasing correlation with ON-state fluorescence as the window widened, underscoring the importance of local structural context in modulating the energetic barriers to translation.

**Figure 3.**
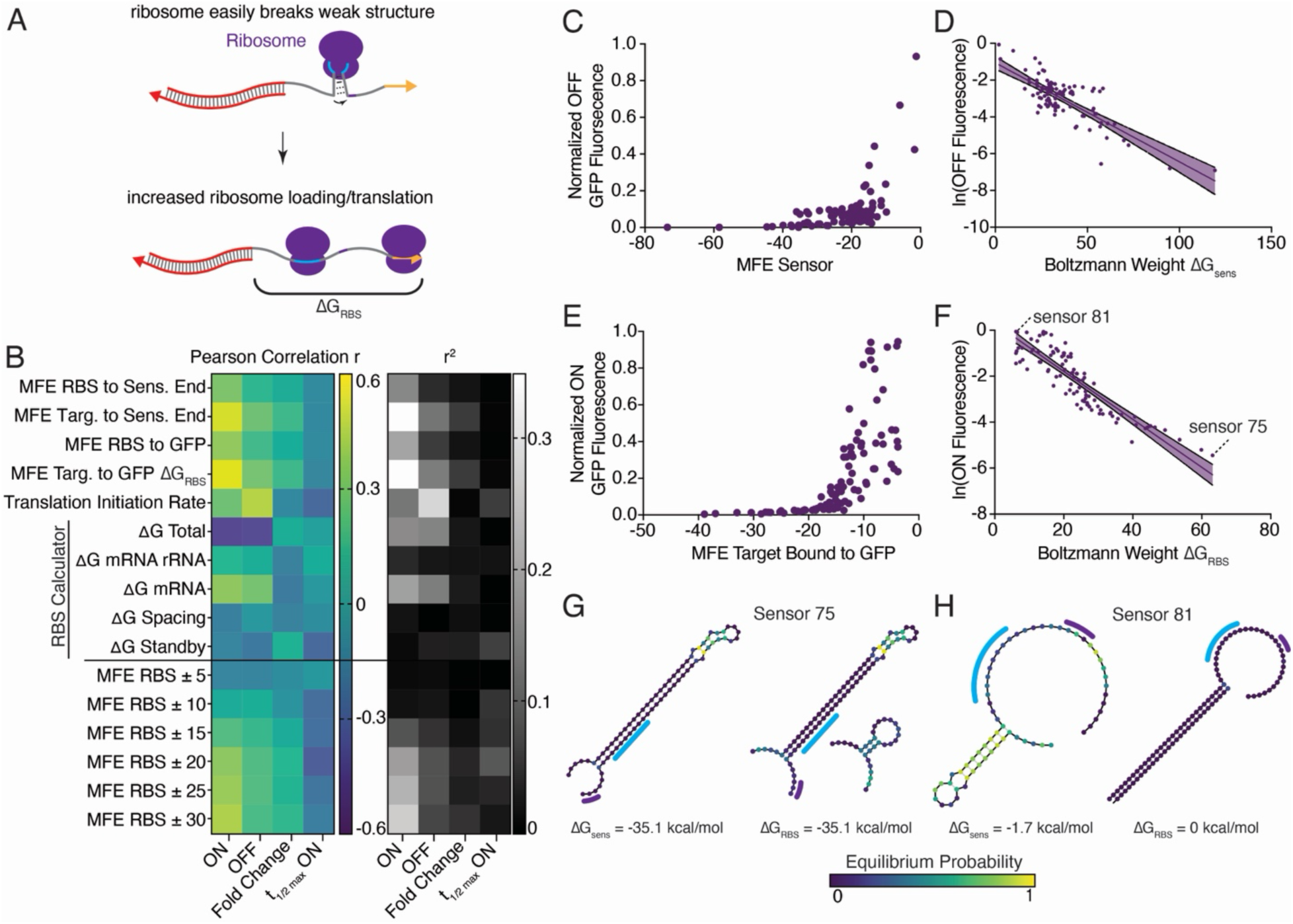
Ribosome accessibility and thermodynamic determinants of translational activation. **(A)** Thermodynamic model of ribosome binding. Structured RBS regions impose energetic penalties on translation initiation that must be overcome for ribosome loading. **(B)** Pearson r correlations between predicted energetic parameters including local MFE windows around the RBS and experimental outputs. **(C)** Relationship between sensor MFE and OFF-state fluorescence, showing stronger repression with lower ΔG_sens_, the free energy of the sensor. **(D)** Log-linear relationship between Boltzmann-weighted structure of the full sensor and fluorescence output, consistent with thermodynamic control of translation initiation (R^2^ =0.61, p<0.0001). **(E)** Relationship between MFE of the target bound-state and ON-state fluorescence. **(F)** Log-linear relationship between the Boltzmann-weighted structure of the RBS region, demonstrating stronger predictive power of structural accessibility metrics (R^2^ =0.78, p<0.0001). **(G)** Representative MFE structure of low-performing sensor 75 wherein the free energy of the structure surrounding the RBS does not change in the presence of target RNA. **(H)** Representative MFE structure of low-performing sensor 81 exhibiting no occlusion of the RBS, resulting in high expression in both ON-and OFF-states. Error bands in D, F represent 95% confidence interval.

The repression of the OFF state was strongly dictated by the intrinsic MFE of the sensor (ΔG_sens_). Sensors with more negative MFE and more stable structure consistently exhibited lower background fluorescence (Figure 3C). Consistent with the model where the rate of translation initiation (*k_init_*) scales exponentially with the free energy of unfolding, if we assume that

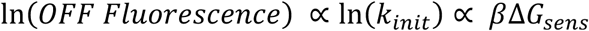

then as the energetic penalty (ΔG_sens_) becomes more negative, the Boltzmann weight decreases, leading to a lower translation rate and therefore lower fluorescence. The negative slope observed in the relationship between OFF fluorescence and the Boltzmann weight of the sensor MFE (Figure 3D) confirms that translation initiation is exponentially suppressed by the energetic cost of unfolding the RBS. In this context, a higher Boltzmann weight indicates a more thermodynamically favorable (lower free energy) structure. Therefore, increased RBS accessibility arises when a greater fraction of the structural ensemble contains the RBS in an unpaired state. As the thermodynamic barrier ΔG_sens_ becomes more negative, the probability of translation initiation events decreases. Across the full library, predicted translation initiation rate correlated with normalized GFP fluorescence (Supplementary Figure 4A), demonstrating agreement between RNA folding energetics and protein expression output.

The accessibility of the RBS in the activated, target-bound state is a more robust predictor of ON-state performance. The MFE of the exposed RBS/start codon sequence after target binding (ΔG_RBS_) correlated strongly with ON fluorescence, and the Boltzmann-weighted RBS accessibility term provided significant predictive power for the induced GFP output (Figure 3E, F). However, ON fluorescence did not correlate strongly with ΔG_sens_ (Supplementary Figure 4B, C), indicating that the thermodynamic barrier encountered by the RBS after target binding influences sensor activation more than the total energy of the sensor. The contrasting energetic profiles of ΔG_sens_ and ΔG_RBS_ are exemplified by two representative poorly performing designs. Sensor 75 exhibits low RBS accessibility in both the OFF (ΔG_sens_ = -35.1 kcal/mol) and ON state (ΔG_RBS_ = -35.1 kcal/mol), in which the target cannot bind and expose the translational machinery (Figure 3G). In contrast, sensor 81 fails to sequester the RBS in the off state (ΔG_sens_ = -1.7 kcal/mol), and the RBS remains exposed in the activated state (Figure 3H). These observations align with the principle that effective repression requires complete overlap between inhibitory structures and the ribosomal footprint, while activation requires significant destabilization of that overlap.

While the observed relationship between fluorescence and Boltzmann weight confirms a thermodynamic dependence, the magnitude of the experimental slope (*β* ≈ 0.1043 mol/kcal) deviates from the previous experimentally measured value of *β* = 0.45 mol/kcal at 37°C^26,45^. The attenuation of the slope suggests translation initiation is not a simple, single-step equilibrium process. The crowdsourced sensors were screened in a transcription-translation (TX-TL) reaction that features an exceptionally high concentration of ribosomes, likely driving initiation kinetics that outpace the RNA’s ability to reach true thermodynamic equilibrium, effectively lowering the sensitivity of the system to structural stability^46,47^. In this high-flux environment, the translation initiation rate may be explained by ribosome loading frequency rather than the global minimum free energy, leading to diminished sensitivity to increases in ΔG_RBS_. Furthermore, non-equilibrium dynamics, such as co-transcriptional folding and the kinetic competition between target RNA binding and RNA degradation likely prevent the sensors from occupying a steady-state Boltzmann distribution^28,48^.

While MFE metrics capture equilibrium structural effects on riboregulator performance, we next asked whether RNA folding kinetics influence translational activation. Since riboregulators fold co-transcriptionally, kinetic trapping or delayed structural rearrangements could modulate RBS accessibility independent of thermodynamic models. If an RNA sensor folds co-transcriptionally into a kinetic trap that exposes the RBS before settling into its thermodynamic MFE, a ribosome can preferentially latch onto it early. To evaluate kinetic effects on the function of crowdsourced sensors, we simulated 100 folding trajectories using Kinfold^49^ and extracted average folding times and early-time energetic features for each RNA sensor, to simulate folding for the riboregulator OFF state and RBS sequestration. We hypothesized that slowly folding RNAs would transiently expose the RBS, enabling higher translation, whereas rapidly folding RNAs would more quickly sequester the RBS and suppress translation (Figure 4A). A representative folding trajectory illustrates the progressive descent from the unfolded state to the MFE structure, with substantial variation in time required to reach the MFE for a single sensor (Supplementary Figure 5A).

**Figure 4.**
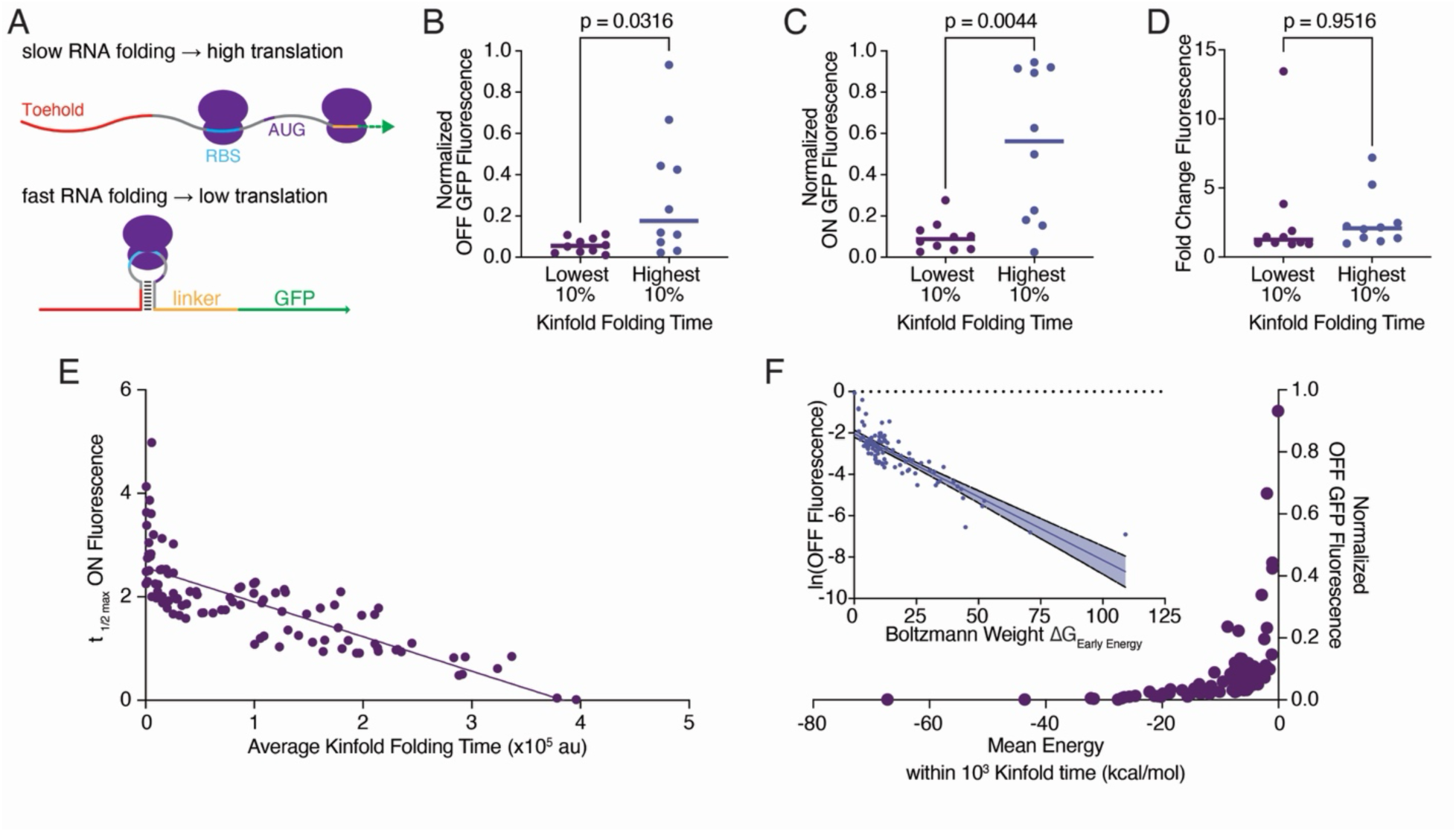
Kinetic determinants of riboregulator function. **(A)** Conceptual model linking RNA folding kinetics to translation. Slow folding delays RBS sequestration and promotes higher translation rates, whereas rapid folding stabilizes inhibitory structures and reduces expression. **(B)** OFF-state fluorescence for the fastest- vs slowest-folding 10% of crowdsourced riboregulators as predicted by Kinfold simulations (p=0.0316, two-tailed t-test). **(C)** ON-state fluorescence stratified by fastest- and slowest- folding extremes (p=0.0044, two-tailed t-test). **(D)** Fold-change activation for fastest- and slowest- folding designs (p=0.9516, two-tailed t-test). **(E)** Relationship between average Kinfold time and time to half-maximal ON fluorescence (t_1/2max_), demonstrating negative correlation between simulated folding time and activation kinetics (R^2^ = 0.61, p<0.0001). **(F)** Mean early-time folding energy (within 10^3^ au Kinfold time units) vs normalized OFF fluorescence. Inset: log-linear relationship between Boltzmann-weighted early folding energies and OFF fluorescence. Error bands represent 95% confidence interval.

Across the crowdsourced library, folding times spanned several orders of magnitude (Supplementary Figure 5B, Supplementary Table 5). Early folding energies also varied considerably (Supplementary Figure 5C), indicating the presence of many intermediate conformational states prior to equilibrium. However, many designs did not immediately fold into their MFE structure. The ratio of mean trajectory energy at 10^3^ au Kinfold time units relative to the MFE ranged widely (Supplementary Figure 5D), and the fraction of trajectories that completed folding within this early time window vary from zero to nearly complete convergence (Supplementary Figure 5E). During early folding, a significant subset of crowdsourced riboregulators remain kinetically partitioned, potentially preserving transient accessibility of translational machinery.

Stratifying designs by folding time revealed significant differences in expression output. The top 10% slowest-folding riboregulators displayed significantly higher OFF-state fluorescence compared to the fastest-folding 10% (Figure 4B), suggesting incomplete or delayed repression during early folding. Similarly, ON-state fluorescence was significantly increased in slow-folding designs (Figure 4C), consistent with enhanced ribosome access. In contrast, fold change did not differ significantly between the kinetic folding time extremes (Figure 4D), indicating that folding kinetics primarily influence expression levels rather than dynamic range. Across the full library, average Kinfold folding time negatively correlated with the time required to reach half-maximal ON fluorescence t_1/2max_ (Figure 4E). Designs with longer simulated folding times activated more rapidly in vitro, consistent with a model in which delayed formation of riboregulator inhibitory structures facilitates earlier ribosome engagement.

We next examined whether early folding energetics predict translational repression. Designs with more negative MFE within the first 10^3^ au Kinfold time exhibited reduced OFF fluorescence (Figure 4F), suggesting that rapid folding of inhibitory conformations enhances repression. Log-linear analysis revealed that OFF fluorescence scales with the Boltzmann-weight associated with early folding states (Figure 4F inset), consistent with thermodynamic control over ribosome accessibility during initial RNA folding. Importantly, the wide variation in early-time energy ratios suggests that many riboregulators do not fully equilibrate early on in folding.

We then sought to test whether slower RNA folding provides a wider kinetic window for ribosome loading. To do so, we selected five crowdsourced sensors that have the same sensor MFE (ΔG_sens_ = -20.1kcal/mol) and compared Kinfold simulations. The distribution of trajectories to reach a fully folded state (Supplementary Figure 6A) revealed a broad spectrum of folding speeds, despite the same equilibrium thermodynamics. Secondary structure analysis (Supplementary Figure 6B) confirmed that while the sensors share a common MFE, they vary slightly in structural features. We then compared the folding times to experimental GFP fluorescence and found a strong positive correlation between average folding time and ON fluorescence, OFF fluorescence, and fold change (Supplementary Figure 6C-E). Sensors that folded more slowly produced higher protein yields, supporting the model that delayed folding activates higher ribosome occupancy. Despite the increase in background leakage, the overall fold change improved as folding time increased, suggesting that kinetic control in conjunction with MFE of the RBS region are dominant factors in determining the dynamic range of riboregulators.

Finally, we examined the energetic states for the five switches early in the folding time (Supplementary figure 6F). The average energy of each trajectory also correlated strongly with OFF-state leakage. Trajectories with higher average energies were associated with increased background GFP fluorescence, indicating that RNAs which remain unfolded for longer are more susceptible to leaky translation initiation in the absence of target RNA. Together, these findings demonstrate that riboregulator performance is governed by both equilibrium stability and co-transcriptional folding dynamics. Slow-folding architectures promote increased translational output, whereas rapid early folding enhances repression. Riboregulator performance is not mediated by energetics alone but is actively regulated by the kinetic competition between RNA secondary structure formation and ribosome binding.

## DISCUSSION

Riboregulator performance is dictated by a coordinated balance between RNA thermodynamics, ribosome accessibility, and folding kinetics. By leveraging community-driven exploration of riboregulator structural space, we uncovered structurally diverse sensors that achieve up to 80-fold activation, demonstrating that high performance is not confined to the canonical toehold switch architecture. The diversity of the solicited riboregulator designs allowed for a robust investigation into the fundamental biophysical principles that distinguish high-performance riboregulator sensors from those that have low dynamic range.

Across the library, translational output was most strongly governed by structural accessibility at the RBS. OFF-state repression scaled with overall sensor stability (ΔG_sens_), whereas ON-state activation depended on the thermodynamic penalty required to expose the RBS (ΔG_RBS_). Fluorescence exhibited an exponential relationship with both free energy terms, consistent with the thermodynamic model of translation initiation. However, the experimentally observed slope was attenuated relative to the theoretical thermodynamic expectation^27^, indicating that translation in the TX-TL system does not operate under simple equilibrium control. Elevated ribosome concentrations likely alter the RNA folding landscape around the RBS prior to full equilibrium, altering the quantitative relationship between predicted RNA folding energetics and the resulting gene expression output.

Beyond equilibrium effects, folding kinetics independently modulated riboregulator behavior. Kinfold simulations revealed significant differences in early-time energy landscapes across sequences with identical MFE, supporting the conclusion that there is no strict relationship between a RNA’s MFE and its folding kinetics. Slowly folding riboregulators exhibited increased OFF-state leakage and elevated ON-state expression, consistent with a model in which delayed inhibitory structure creates transient windows of ribosome accessibility. Early-time energetic metrics correlated with OFF-state fluorescence, underscoring that folding trajectories, not just endpoint MFE, shapes translational output and riboregulator function. While kinetics influenced expression levels in the absence and presence of target, dynamic range was dictated primarily from a balance between repression strength and RBS accessibility. Riboregulator variants with identical MFE displayed different expression outputs due to differences in folding dynamics, demonstrating that equilibrium stability sets a baseline repression threshold, whereas kinetic accessibility determines how effectively that threshold is reached.

Although the crowdsourced riboregulator approach expanded structural diversity, the library still represents a sparse sampling of the massive riboregulator design space^6^, limiting our ability to generalize quantitative relationships across all architectures. While highly controlled and reproducible, all measurements were performed in a cell-free TX-TL system, which does not fully recapitulate the molecular crowding, RNA degradation pathways, transcriptional dynamics, and ribosome competition in living cells^50^. Despite our analyses relying on equilibrium folding models and stochastic kinetic simulations that approximate co-transcriptional folding and ribosome-RNA coupling during translation initiation, our findings support a multi-parameter framework for riboregulator design: thermodynamic stability governs repression, RBS accessibility dictates activation, and folding kinetics modulate expression magnitude. More broadly, this work demonstrates that expanding exploration of RNA structure space enabled through crowdsourced design can reveal non-intuitive structural solutions and refine our understanding of programmable RNA control of gene expression.

## METHODS

### Crowdsourced riboregulator design

Participants were provided viral genome fragments from the SARS-CoV-2 nucleocapsid (N) gene, Zika Virus capsid/precursor membrane/envelope (CME) genes, Human Rotavirus outer capsid VP4 gene, and Influenza A hemagglutinin (HA) gene (Figure 1A, Supplementary Table 6). No explicit performance guidance was provided to the community, but participants were instructed to design riboregulators that minimized basal translation in the absence of target RNA and maximized translation upon target RNA binding. The community was allowed to use any available RNA folding software, including ViennaRNA^49^ and NUPACK^51,52^. Strand orientation was unfixed—designers could use either the sense or antisense of target genes—and participants were asked to model RNA folding at 37°C. Designs were constrained to be less than 200 nucleotides, feature the RBS sequence ‘AGAGGAGA’ and the 21-nucleotide linker sequence ‘AACCUGGCGGCAGCGCAAAAG’ between the riboregulator and mut3bGFP-asv reporter. The RBS and linker sequences were held constant to isolate regulatory effects arising solely from riboregulator structural differences and rule out RBS-driven effects on translation initiation. No additional structural or energetic filters were imposed prior to experimental testing. Designs were not required to fold in any structure to be submitted by participants. All submissions were filtered using an automated script for in-frame stop codons and to verify that the sequence length between the start codon and fixed linker was a multiple of three, ensuring that the downstream mut3bGFP-asv reporter remained in-frame. Designs containing premature stop codons or incorrect reading frame lengths were returned to participants for revision.

### Plasmids and oligonucleotides

Control toehold switches were designed with NUPACK 4.0.0 using the toehold-VISTA code^31^. All crowdsourced and control riboregulators were ordered with a common T7 promoter and linker sequence (Supplementary Table 6). Target RNAs were ordered with a 5’ T7 promoter and common 3’ sequence for amplification (Supplementary Table 6). All riboregulators and targets were ordered as ssDNA Ultramers from IDT. Sequences for all primers used in this study can be found in Supplementary Table 6. All oligonucleotides were purchased from IDT.

### Cell-free transcription-translation template preparation and characterization

Riboregulator oligos were appended to the mut3bGFP-asv reporter for transcription-translation (TX-TL) by fusion PCR. ssDNA ultramers were used as a forward primer to amplify a linear mut3bGFP-asv fragment with T7 terminator using Q5 High-Fidelity 2x Master Mix (New England Biolabs M0492L). Following verification by gel electrophoresis, riboregulator reporter amplicons were purified (New England Biolabs T1130L) and normalized to 50 nM. Target DNA was prepared by amplification of ssDNA ultramers using a T7 forward primer and a common reverse primer using Q5 2x Master Mix. Following purification, target DNA template was normalized to 200 nM. Riboregulator characterization was performed in cell-free TX-TL reactions using New England Biolabs PURExpress *In Vitro* Protein Synthesis Kit (E6800S). 1.5 μL of riboregulator and target DNA templates were added to each 5 μL cell-free reaction condition with 0.08 μL RNase Inhibitor (New England Biolabs M0314S). Each riboregulator design was tested either with cognate target (ON state) or with non-cognate decoy target RNA (OFF state). Reactions were sealed, spun down, and placed in a BioTek Cytation 5 multimode reader (Agilent) pre-warmed to 37**°**C. After an initial shaking at 1200 rpm for 30 seconds to ensure adequate mixing, measurements were taken every 45 seconds at 485/20 emission 528/20 from the bottom of the plate with a gain of 85 for 6 hours.

### Data normalization and calculation of fold change

Fluorescence data were first smoothed to reduce noise in measurements using a moving five-point average of each time point and the preceding and following data^14^. Background signal was calculated using averaged intensities of blank reactions that received cell-free extract alone (*F_min_*), and the maximal fluorescence value was calculated using the averaged intensities of three reactions with T7-driven expression of the mut3bGFP construct with no inhibitory structures in the 5’ UTR (*F_max_*). The minimum value of each well was then adjusted to zero. Data normalization to positive and negative control fluorescence was used to scale experimental data and account for variability in experiments conducted on different days. Fluorescence normalization for each sample followed the formula:

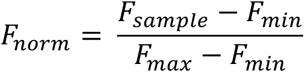

Fold change calculations for fluorescence data were done by dividing the average signal from wells with cognate target by the non-cognate decoy target wells for each riboregulator. For rate fold change, the difference in the rate of GFP fluorescence increase between cognate and non-cognate target wells was taken. The rate of change was calculated using the slope. At the two-hour time point, the rate reported was calculated by:

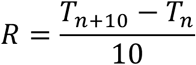

where *T_n_* is the normalized data at the two-hour time point and *R* is the rate reported for *T_n_* in AU-min^-1^. Fold change rate was then calculated as with the fluorescence data. The values reported in Supplementary Table 1 represent the mean of n=3 biological replicates gathered in three separate experiments on different days

### Calculations made with ViennaRNA, NUPACK, and RBS Calculator

RNA secondary structure predictions were performed using the ViennaRNA package. Minimum free energy (MFE) structures were calculated under default parameters at 37**°**C. For each riboregulator sequence, we computed MFE structure and corresponding free energy of the sensor, target, and sensor-target complex (Figure 2). For more granular MFE calculations around the RBS, a custom script was written to identify the RBS/start index in each sensor and the last index of the target binding domain (Supplementary Figure 3). Each sensor sequence was indexed using these positions to model sub-sequence MFEs from the target binding domain to the end of the sensor (not including the linker), or up to the start of the mut3bGFP-asv (including the linker). Window calculations of MFE (Figure 3B) were conducted with all available nucleotides if the sampled window surpassed the available length preceding or following the RBS. Independent thermodynamic validation and ensemble analyses were performed using NUPACK 4.0.0 and compared with ViennaRNA-derived pairing probabilities to ensure consistency across modeling frameworks.

The kinetic accessibility of MFE metrics and the presence of transient intermediates were evaluated using the Kinfold extension of ViennaRNA, which implements a Monte Carlo algorithm to simulate stochastic folding trajectories. For each sequence, 100 independent folding trajectories were simulated. Each trajectory was allowed to proceed for a maximum of 10^6^ arbitrary Kinfold time units. The starting configuration for all simulations was the unfolded riboregulator source sequence, and the rates of all possible nucleotide pairing and unpairing transitions were then computed. For each riboregulator sequence, the fraction of trajectories that reached the target MFE structure (ΔG_sens_) within the simulation time limit was calculated. To assess the stability of folding intermediates and early folding, the mean energy was calculated during the early folding phase, defined as the first 10^3^ arbitrary Kinfold time units. An energy ratio was derived by comparing the mean early folding energy to the final MFE energy (ΔG_early_/ΔG_sens_) to quantify the energetic distance from equilibrium state during the initial folding window. Full trajectory traces were recorded for the first 10 trajectories of each variant for detailed analysis (Supplementary Figure 5-6).

Translation initiation rates were estimated using the RBS Calculator^26–29,53^. For each riboregulator sequence, the 5’ untranslated region (from the transcriptional start site, TSS) through the 200^th^ index after the TSS was submitted to the model. The RBS calculator estimates translation initiation rate (TIR) based on a thermodynamic framework that accounts for mRNA:rRNA hybridization free energy, start codon interaction energy, spacing penalties between Shine-Dalgarno sequence and the start codon, mRNA secondary structure unfolding free energy, and standby site accessibility. For each crowdsourced riboregulator design, the calculated position for the common RBS was used to index the RBS Calculator output file and save the corresponding energy parameters (Figure 3, Supplementary Table 4).

### Consensus Structure Calculation

For each sensor sequence within a designated cohort (for example, highest/lowest fold change), the thermodynamic partition function *Z* was computed using the ViennaRNA package. The partition function represents the Boltzmann—weighted sum of all possible secondary structures *σ* for a given sequence:

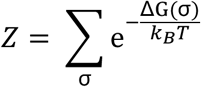

From this ensemble, we derived the base-pairing probability (BPP) matrix, where each element *P_ij_*: denotes the probability of a base pair forming between residues *i* and *j*. To capture the collective behavior of the population, a mean cohort BPP matrix (*̄P^ij^*) was calculate by averaging the individual matrices across all *N* sequences in the cohort:

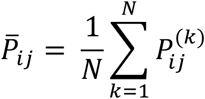

To provide a physical frame of reference for the ensemble data, we generated a consensus structural scaffold using a positional majority-rule approach across the MFE structures aligned based on the common RBS motif AGAGGAGA to account for riboregulator sequences of varying lengths. At each position *j* within the MFE structure, the ensemble pairing probability of each nucleotide was then calculated as the marginal sum of its pairing partners within the window:

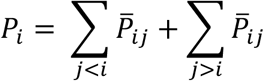

This *P_i_* metric was mapped onto the radial consensus scaffold, providing a visual proxy for the thermodynamic accessibility of the RBS and its surrounding regulatory secondary structure.

## ASSOCIATED CONTENT

### Supplementary Information

Supplementary information includes additional experimental data and tables for DNA sequences used in the study (PDF).

## Author Contributions

J.M.R. and A.A.G. designed the study. J.M.R., G.M., and D.F. carried out experimental work. J.M.R. and G.M. conducted data analysis. A.A.G. supervised the study and acquired funding. J.M.R., G.M., and A.A.G wrote the manuscript. All authors edited the manuscript.

## ACKNOWLEDGMENTS

This work was supported by startup funds from Boston University; a National Institutes of Health (NIH) U01 award (1U01AI148319), R01 award (1R01EB031893), and a NIH Director’s Transformative Research Award (R01EB037112) to A.A.G. J.M.R. was supported by the NIH Training Program in Synthetic Biology and Biotechnology (1T32GM130546) and a National Science Foundation Graduate Research Fellowship (2234657). The content is solely the responsibility of the authors and does not necessarily represent the official views of the National Institutes of Health or the National Science Foundation.

## REFERENCES

1. Breaker, R. R. Riboswitches and the RNA World. Cold Spring Harb Perspect Biol 4, a003566 (2012).

2. Serganov, A. & Nudler, E. A decade of riboswitches. Cell 152, 17–24 (2013).

3. Lucks, J. B., Qi, L., Mutalik, V. K., Wang, D. & Arkin, A. P. Versatile RNA-sensing transcriptional regulators for engineering genetic networks. Proceedings of the National Academy of Sciences 108, 8617–8622 (2011).

4. Chappell, J., Takahashi, M. K. & Lucks, J. B. Creating small transcription activating RNAs. Nat Chem Biol 11, 214–220 (2015).

5. Isaacs, F. J. et al. Engineered riboregulators enable post-transcriptional control of gene expression. Nat Biotechnol 22, 841–847 (2004).

6. Green, A. A., Silver, P. A., Collins, J. J. & Yin, P. Toehold Switches: De-Novo-Designed Regulators of Gene Expression. Cell 159, 925–939 (2014).

7. Zhang, Q. et al. Predictable control of RNA lifetime using engineered degradation-tuning RNAs. Nat Chem Biol 17, 828–836 (2021).

8. Gao, Y. et al. Programmable trans-splicing riboregulators for complex cellular logic computation. Nat Chem Biol 21, 758–766 (2025).

9. Mandal, M. & Breaker, R. R. Gene regulation by riboswitches. Nat Rev Mol Cell Biol 5, 451–463 (2004).

10. Gottesman, S. et al. Small RNA regulators and the bacterial response to stress. Cold Spring Harb Symp Quant Biol 71, 1–11 (2006).

11. Vigouroux, A. & Bikard, D. CRISPR Tools To Control Gene Expression in Bacteria. Microbiol Mol Biol Rev 84, e00077–19 (2020).

12. Kim, J. et al. De novo-designed translation-repressing riboregulators for multi-input cellular logic. Nat Chem Biol 15, 1173–1182 (2019).

13. Ma, D. et al. Multi-arm RNA junctions encoding molecular logic unconstrained by input sequence for versatile cell-free diagnostics. *Nat*. Biomed. Eng 6, 298–309 (2022).

14. Pardee, K. et al. Rapid, Low-Cost Detection of Zika Virus Using Programmable Biomolecular Components. Cell 165, 1255–1266 (2016).

15. Thavarajah, W. et al. Point-of-Use Detection of Environmental Fluoride via a Cell-Free Riboswitch-Based Biosensor. ACS Synth. Biol. 9, 10–18 (2020).

16. Karlikow, M. et al. Field validation of the performance of paper-based tests for the detection of the Zika and chikungunya viruses in serum samples. *Nat*. Biomed. Eng 6, 246–256 (2022).

17. Carr, A. R. et al. Toward Mail-in-Sensors for SARS-CoV-2 Detection: Interfacing Gel Switch Resonators with Cell-Free Toehold Switches. ACS Sens. 7, 806–815 (2022).

18. Green, A. A. et al. Complex cellular logic computation using ribocomputing devices. Nature 548, 117–121 (2017).

19. Zhang, M. et al. High-resolution and programmable RNA-IN and RNA-OUT genetic circuit in living mammalian cells. Nat Commun 15, 8768 (2024).

20. Kaseniit, K. E. et al. Modular, programmable RNA sensing using ADAR editing in living cells. Nat Biotechnol 41, 482–487 (2023).

21. Kaseniit, K. E. et al. Modular, programmable RNA sensing using ADAR editing in living cells. Nat Biotechnol 41, 482–487 (2023).

22. Amalfitano, E. et al. A glucose meter interface for point-of-care gene circuit-based diagnostics. Nat Commun 12, 724 (2021).

23. Sadat Mousavi, P., et al. A multiplexed, electrochemical interface for gene-circuit-based sensors. Nat. Chem. 12, 48–55 (2020).

24. Pardee, K. et al. Paper-Based Synthetic Gene Networks. Cell 159, 940–954 (2014).

25. Li, Y. et al. Bispecific Coiled-Coil Reporters Enable Rapid, Multiplexed Detection of Mycoplasma via Toehold Switch Sensors. 2026.05.28.728408 Preprint at 10.64898/2026.05.28.728408 (2026).

26. Salis, H. M., Mirsky, E. A. & Voigt, C. A. Automated design of synthetic ribosome binding sites to control protein expression. Nat Biotechnol 27, 946–950 (2009).

27. Espah Borujeni, A., Channarasappa, A. S. & Salis, H. M. Translation rate is controlled by coupled trade-offs between site accessibility, selective RNA unfolding and sliding at upstream standby sites. Nucleic Acids Research 42, 2646–2659 (2014).

28. Espah Borujeni, A. & Salis, H. M. Translation Initiation is Controlled by RNA Folding Kinetics via a Ribosome Drafting Mechanism. J. Am. Chem. Soc. 138, 7016–7023 (2016).

29. Espah Borujeni, A., et al. Precise quantification of translation inhibition by mRNA structures that overlap with the ribosomal footprint in N-terminal coding sequences. Nucleic Acids Res 45, 5437–5448 (2017).

30. Mustoe, A. M. et al. Pervasive Regulatory Functions of mRNA Structure Revealed by High-Resolution SHAPE Probing. Cell 173, 181–195.e18 (2018).

31. Robson, J. M. & Green, A. A. Toehold-VISTA: a machine learning approach to decipher programmable RNA sensor-target interactions. Nucleic Acids Res 54, gkag097 (2026).

32. Angenent-Mari, N. M., Garruss, A. S., Soenksen, L. R., Church, G. & Collins, J. J. A deep learning approach to programmable RNA switches. Nat Commun 11, 5057 (2020).

33. Valeri, J. A. et al. Sequence-to-function deep learning frameworks for engineered riboregulators. Nat Commun 11, 5058 (2020).

34. Riley, A. T., Robson, J. M., Ulanova, A. & Green, A. A. Generative and predictive neural networks for the design of functional RNA molecules. Nat Commun 16, 4155 (2025).

35. Riley, A. T. et al. A Unified Framework for Model-Informed and Agentic RNA Design. 2025.06.17.659751 Preprint at 10.1101/2025.06.17.659751 (2026).

36. Andreasson, J. O. L. et al. Crowdsourced RNA design discovers diverse, reversible, efficient, self-contained molecular switches. Proc Natl Acad Sci U S A 119, e2112979119 (2022).

37. Robson, J. M. & Green, A. A. Closing the loop on crowdsourced science. Proceedings of the National Academy of Sciences 119, e2205897119 (2022).

38. Wayment-Steele, H. K. et al. Deep learning models for predicting RNA degradation via dual crowdsourcing. Nat Mach Intell 4, 1174–1184 (2022).

39. He, S., et al. Ribonanza: deep learning of RNA structure through dual crowdsourcing. bioRxiv 2024.02.24.581671 (2024) doi:10.1101/2024.02.24.581671.

40. Choe, C. A. et al. Compact RNA sensors for increasingly complex functions of multiple inputs. Nat. Chem. 17, 1839–1852 (2025).

41. Tangpradabkul, T. et al. Minimization of the E. coli ribosome, aided and optimized by community science. Nucleic Acids Res 52, 1027–1042 (2024).

42. Koepnick, B. et al. De novo protein design by citizen scientists. Nature 570, 390–394 (2019).

43. Lee, J. et al. RNA design rules from a massive open laboratory. Proceedings of the National Academy of Sciences 111, 2122–2127 (2014).

44. Voelz, V. A., Pande, V. S. & Bowman, G. R. Folding@home: Achievements from over 20 years of citizen science herald the exascale era. Biophys J 122, 2852–2863 (2023).

45. Hao, Y. et al. Quantifying the sequence-function relation in gene silencing by bacterial small RNAs. Proc Natl Acad Sci U S A 108, 12473–12478 (2011).

46. Vezeau, G. E. & Salis, H. M. Tuning Cell-Free Composition Controls the Time Delay, Dynamics, and Productivity of TX-TL Expression. ACS Synth. Biol. 10, 2508–2519 (2021).

47. Karzbrun, E., Shin, J., Bar-Ziv, R. H. & Noireaux, V. Coarse-Grained Dynamics of Protein Synthesis in a Cell-Free System. Phys. Rev. Lett. 106, 048104 (2011).

48. Mustoe, A. M., Corley, M., Laederach, A. & Weeks, K. M. mRNA structure regulates translation initiation: a mechanism exploited from bacteria to humans. Biochemistry 57, 3537–3539 (2018).

49. Lorenz, R. et al. ViennaRNA Package 2.0. Algorithms for Molecular Biology 6, 26 (2011).

50. Takahashi, M. K. et al. Rapidly Characterizing the Fast Dynamics of RNA Genetic Circuitry with Cell-Free Transcription–Translation (TX-TL) Systems. ACS Synth Biol 4, 503–515 (2015).

51. Zadeh, J. N. et al. NUPACK: Analysis and design of nucleic acid systems. Journal of Computational Chemistry 32, 170–173 (2011).

52. Fornace, M. E. et al. NUPACK: Computational Nucleic Acid Analysis and Design. ACS Synth. Biol. 15, 1426–1441 (2026).

53. Cetnar, D. P. & Salis, H. M. Systematic Quantification of Sequence and Structural Determinants Controlling mRNA stability in Bacterial Operons. ACS Synth. Biol. 10, 318–332 (2021).

