## Supplementary Information for "Crowdsourced riboregulators reveal design principles for programmable RNA switching"

### Supplementary Information: Crowdsourced riboregulator design reveals principles for programmable RNA switching

This PDF file includes:

Supplementary Figures 1 to 6

Additional Supplementary Tables:

Supplementary Table 1: Crowdsourced riboregulator sequences with experimental measurements

Supplementary Table 2: Control toehold-VISTA riboregulators with associated experimental measurements

Supplementary Table 3: Calculated structural and energetic features of crowdsourced riboregulator library

Supplementary Table 4: Calculated biophysical parameters relating to RBS accessibility metrics

Supplementary Table 5: Kinfold trajectory energy and time calculations

Supplementary Table 6: Sequences for target sources and template construction

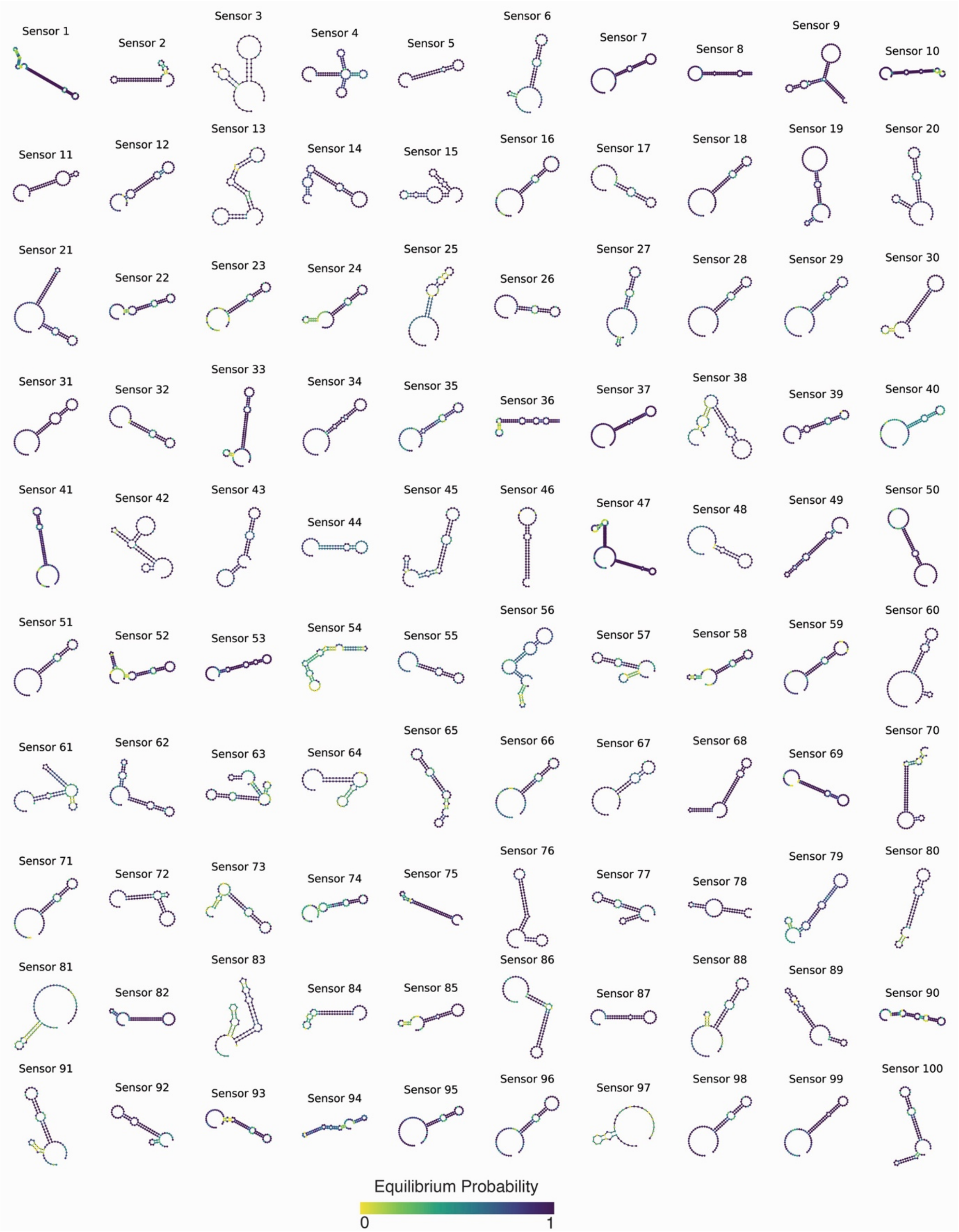


**Supplementary Figure 1. Secondary structure ensembles of community-designed riboregulators.** Each structure represents a unique architectural solution submitted by the community to regulate ribosome binding. Color scale indicates equilibrium probability of each nucleotide bound, ranging from 0 to 1.

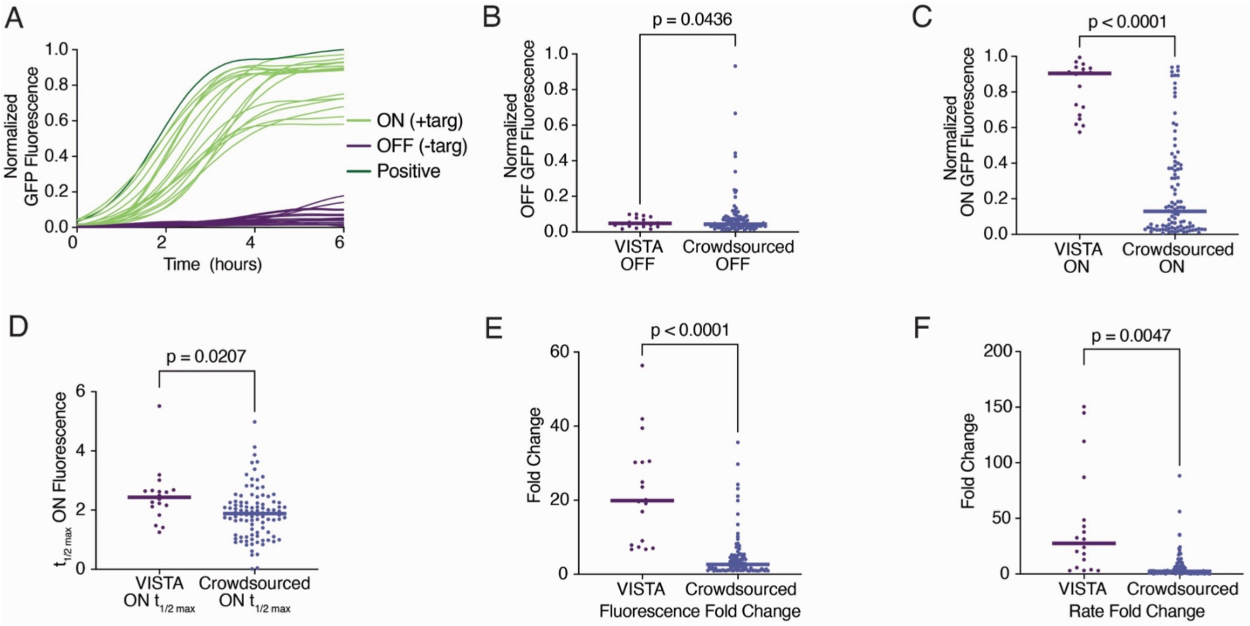


**Supplementary Figure 2. Comparative analysis of crowdsourced and canonical toehold-VISTA sensors activation dynamics and fold change. (A)** Kinetic traces of normalized GFP fluorescence over 6 hours in the presence of cognate (ON) or non-cognate decoy (OFF) RNA targets. **(B)** Distribution of OFF-state fluorescence values between VISTA-designed toehold switches and crowdsourced sensors. **(C)** Comparison of ON-state fluorescence activation between VISTA and crowdsourced sensors. **(D)** Time-to-activation comparison ($t_{1/2max}$) for VISTA and crowdsourced sensors. **(E)** Fold change in fluorescence and **(F)** fold change in rate distributions for VISTA and crowdsourced libraries, demonstrating that select crowdsourced designs significantly outperform standard toehold switch sensors.

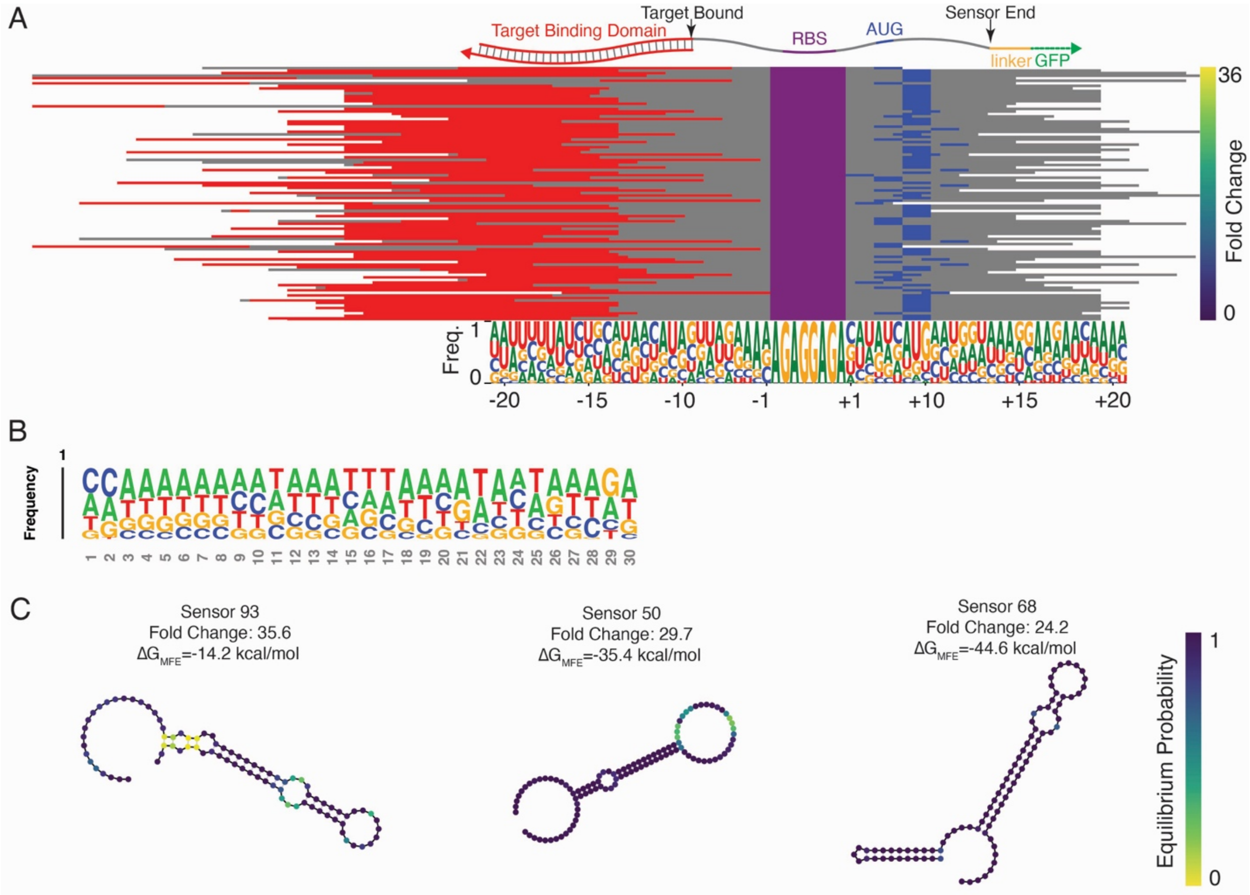


**Supplementary Figure 3. Structural and sequence motifs governing crowdsourced riboregulator performance. (A)** Alignment of sensor architectures relative to the RBS, ranked by fold change. All target binding domains remain at the 5' end of the RBS, with varying spacing between the target bound position, RBS, and start codon. Sensors also vary widely in spacing between the start codon and the linker. **(B)** Sequence logo displaying nucleotide frequency for the first 30 nucleotides of all targets. Targets shorter than 30 nucleotides were padded for proper alignment. **(C)** MFE structures for three top-performing sensors, showing diverse structures for successful RBS sequestration.

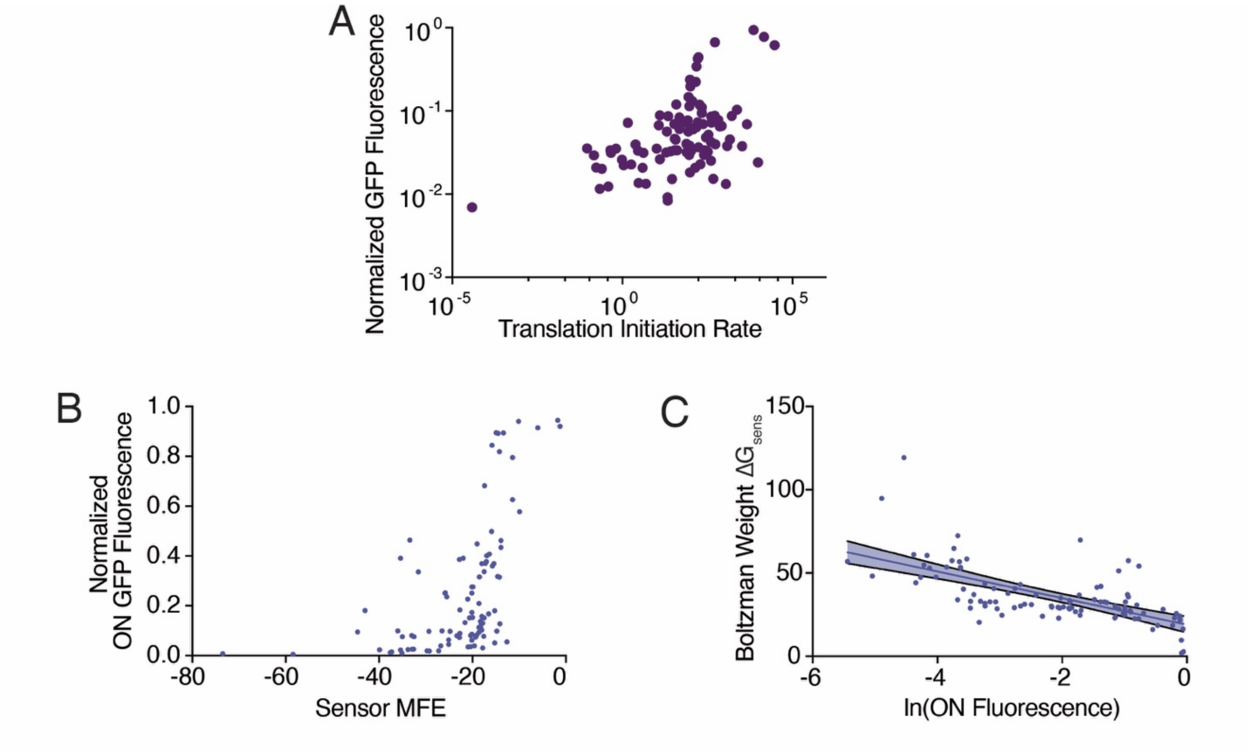


**Supplementary Figure 4. RNA folding energetics and experimental fluorescence. (A)** Normalized GFP fluorescence as a function of predicted initiation rate. **(B)** Relationship between sensor MFE and ON-state fluorescence, showing a sharp increase in performance at specific energetic thresholds. **(C)** Boltzmann-weighted ensemble energy for $\Delta G_{sens}$ correlates poorly with the natural log of ON fluorescence ($R^2$ =0.42, p<0.0001). Error bands represent 95% confidence interval.

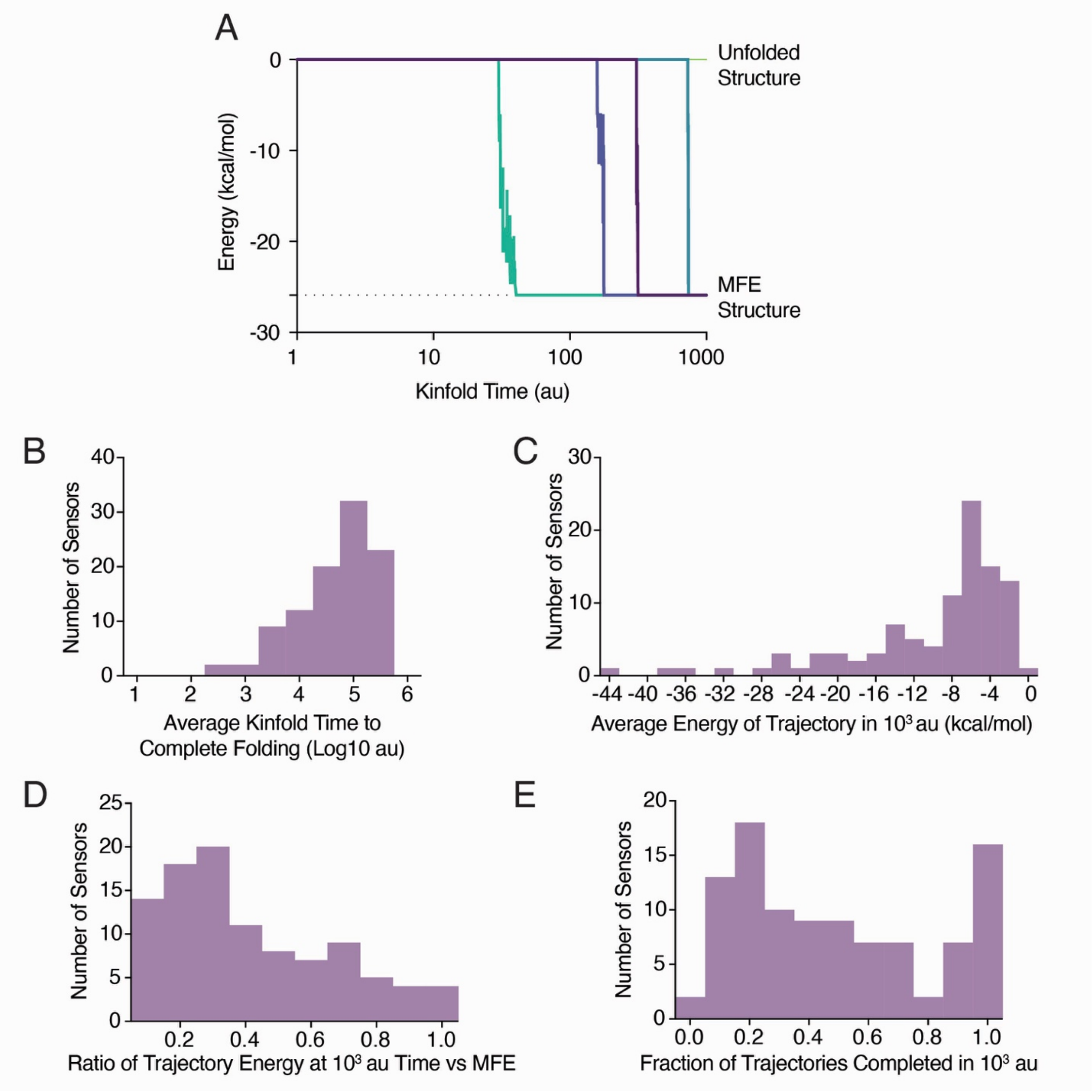


**Supplementary Figure 5. Characterization of RNA folding landscape using Kinfold.** Folding trajectories were run using ViennaRNA's Kinfold package for each crowdsourced sensor. **(A)** Five representative Kinfold trajectories for sensor 95 showing the time-dependent transition from an unfolded state to the MFE structure. While four sensors were able to reach the MFE in $10^3$ arbitrary Kinfold time units, one did not. **(B)** Histogram representing the time for each sensor to complete folding (average of n=100 trajectories for each sensor). Most trajectories took longer than $10^3$ au, and $10^6$ was the longest allowed trajectory time. **(C)** Representative free energy reached in the first $10^3$ au, representing $\Delta G_{early}$ **(D)** The ratio of each sensor's average energy over 100 trajectories in comparison to the MFE for that sensor. **(E)** Fraction of trajectories that complete folding of the MFE structure in $10^3$ au.

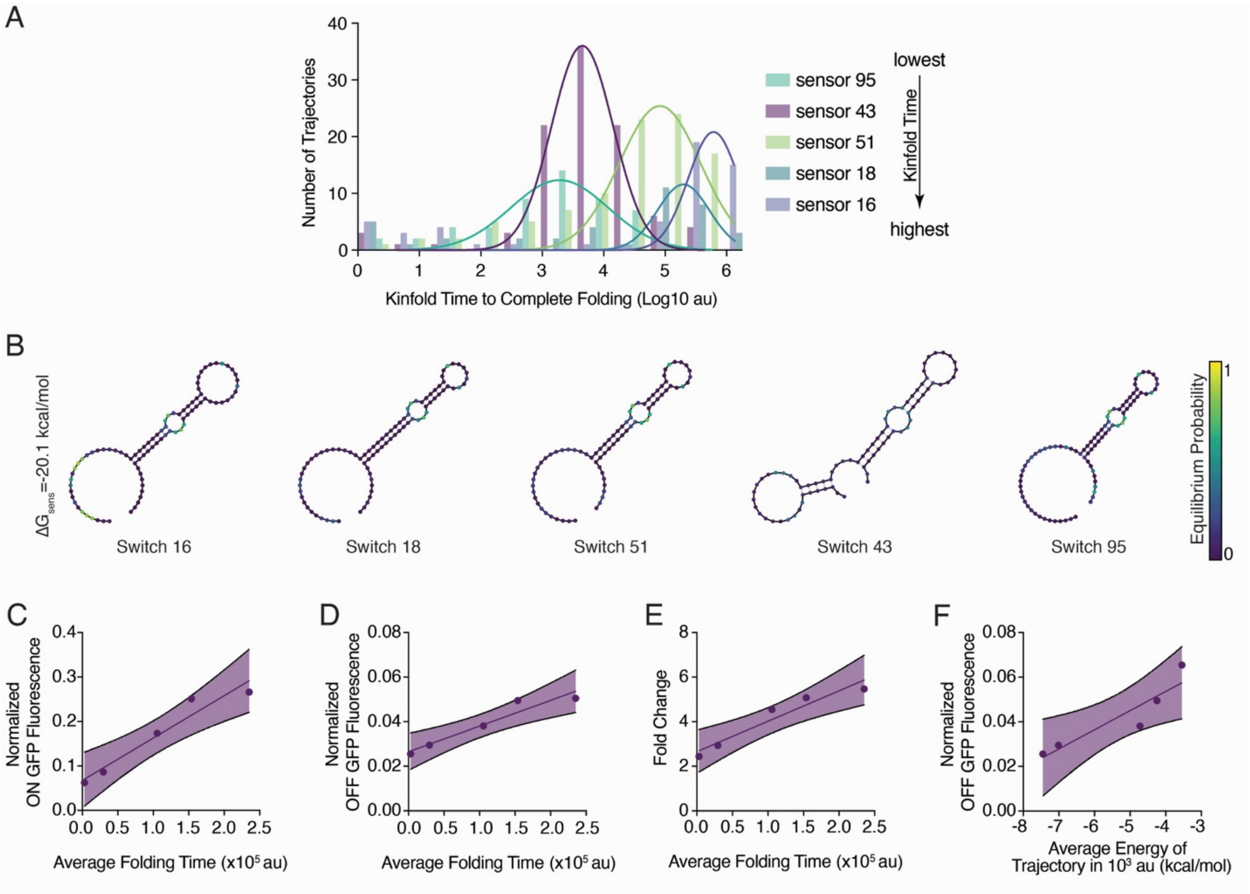


**Supplementary Figure 6. Impact of folding kinetics on sensor performance.** For five crowdsourced designs that feature the same MFE, 100 trajectories were run for a maximum time of $10^6$ arbitrary Kinfold time units. **(A)** Representative histograms for the Kinfold time required to reach the MFE structure are overlayed with a gaussian fit of the distribution. (sensor 43 $R^2$ =0.98, sensor 16 $R^2$ =0.86, sensor 18 $R^2$ =0.47, sensor 95 $R^2$ =0.90, sensor 51 $R^2$ =0.87). **(B)** MFE structures for the five representative crowdsourced sensors with varying folding speeds, ranked from highest to lowest average Kinfold time to reach MFE structure. Correlation between average folding time per trajectory and **(C)** ON-state GFP fluorescence ($R^2$ =0.95, p=0.0044), **(D)** OFF-state GFP fluorescence ($R^2$ =0.92, p=0.0094), **(E)** fold change ($R^2$ =0.92, p=0.0092), and **(F)** average early energy $\Delta G_{early}$ ($R^2$ =0.84, p=0.0267), suggesting that sensors with longer folding times exhibit higher basal leakage and varied ON-state dynamics. Error bands represent 95% confidence interval.
